## Supplemental Table 1 for "BCAS2 promotes primitive hematopoiesis by sequestering β-catenin within the nucleus"

**Primers Used for Genotyping**

| **Strain** | **Directions** | **Sequence (5' to 3')** |
|---|---|---|
| *Flk1*-Cre mouse | Forward | CGGTTATTCAACTTGCACCAC |
| | Reverse | CAGGACTGAAAGCCCAGACT |
| *Bcas2*$^{Flox/Flox}$ mouse | Forward | ATTCCAGCAGTTGGTGTGGG |
| | Reverse | CATTGCTGGACAGAAGGTGAG |
| *Flk1*-Cre;*Bcas2*$^{Flox/Flox}$ mouse | Forward | AGGTGTATGAATGCCTGAACAAG |
| | Reverse | CATTGCTGGACAGAAGGTGAG |
| *bcas2* knockout zebrafish | Forward | TGCACATACAGTAATAGGCTTACCC |
| | Reverse | GTCTGATTTGCATCAAAAGATGTGA |
