## Supplemental Table 2 for "BCAS2 promotes primitive hematopoiesis by sequestering β-catenin within the nucleus"

**Primers Used for Reverse Transcription-PCR**

| **Symbol** | **Directions** | **Sequence (5’ to 3’)** |
|---|---|---|
| *ddb2* | Forward | CGCTCACTTAAATCTTACAAGCTGC |
| | Reverse | AGTCTGGGGTACGAGACAATATCT |
| | Reverse | TTGGCAAAGACTTGTATAACGGATC |
| *acox* | Forward | ATCCAGATTTCCAACATGAAGACCT |
| | Reverse | AGGTGCCTAACATATGGAATGACTC |
| | Forward | TTCTCTCTCGAAGTGAGCGATATGA |
| | Reverse | TCTGTGCGTAGGTGCCAATAATCTC |
| *ubap2l* | Forward | GAATGTTGGTGTCAATGCATCAGCT |
| | Reverse | CTGCGTCTTTGTGTAAACAGAGCCT |
| *cdk16* | Forward | ATGAGGAAGATCAAACGGCAGCTGT |
| | Reverse | AAATTTTGCGACCGGGGTTGTTTCT |
| *rbbp4* | Forward | TGGGACACTCGTTCCAATAACACAT |
| | Reverse | CTGGAAGATTTCATCTTTGTGCGAC |
| *her6* | Forward | GGAGAAAAGAAGAAGAGCGAGAATC |
| | Reverse | GCTGCATGTTTCTGAGATGTTTCAC |
| *cirbpb* | Forward | TCCTACAGAGACGGTTACGACAGTT |
| | Reverse | CCTCACCAGGCTTGAAACTCTTACA |
| *cdkn2aip* | Forward | TCGCTAATGAGGACTTGTCTTTGGA |
| | Reverse | ATTTATCTTGTCCATCACACGCTGC |
| *mapk1* | Forward | GTCTGTTGGTTGTATTCTGGCTGAG |
| | Reverse | AGGATCATAGTACTGCTCCAGGTAC |
| *add1* | Forward | AGGATGCTGGATAATCTGGGCTACA |
| | Reverse | GTTGCGCATCTCTTGGACTTCTTTC |
| *Mdm4*-FL | Forward | GGTCAGGTGTCCAGTGAGTCAATAA |
| | Reverse | AAACCATCTGAGGAGTCTTCATCTG |
| *Mdm4*-S | Forward | GAAGATCCTGGTCAGACTCCTCAGA |
| | Reverse | CGGGAGAGAGTTGATTGGTGTGAAT |
| *β-catenin* | Forward | GATGACGATGTGGATAATCAGGTGC |
| | Reverse | GAACTGTGTGGAAGGTATCTGCATG |
| | Forward | GGAGACAATTTCTCATTATTCTGGC |
| | Reverse | TCAGTAGTTTGGTGAGCTCTGGAAT |
| *β-actin* | Forward | ATGGATGATGAAATTGCCGCAC |
| | Reverse | ACCATCACCAGAGTCCATCACG |
