## Supplemental Figures for "BCAS2 promotes primitive hematopoiesis by sequestering β-catenin within the nucleus"

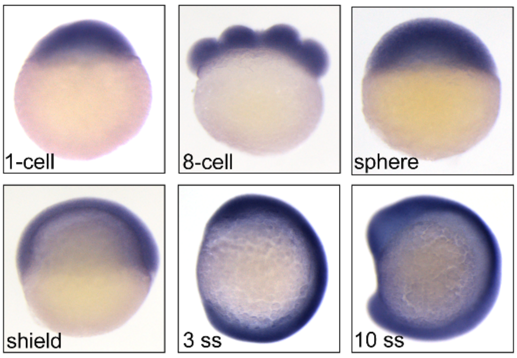


**Figure S1. Expression patterns of *bcas2* in wild-type embryos during development.**

Expression of *bcas2* in wild-type embryos at the indicated developmental stages was analyzed using whole-mount *in situ* hybridization.

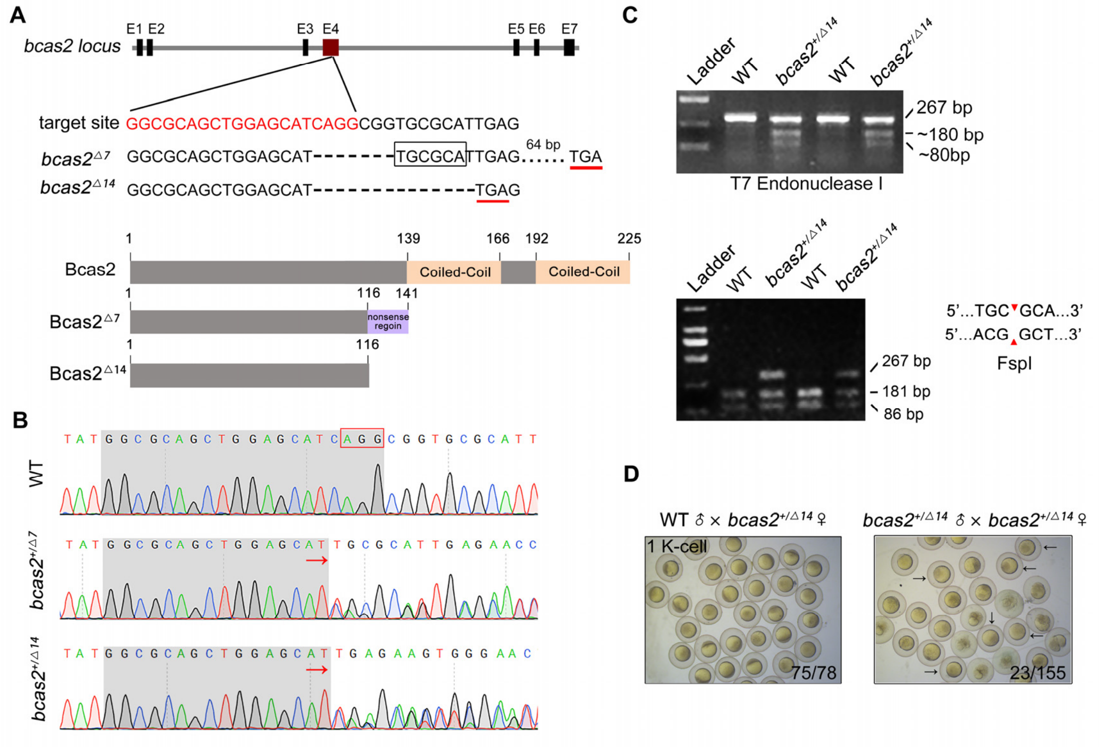


**Figure S2. Zebrafish *bcas2* mutants are generated by using CRISPR/Cas9 system.**

(A) Schematic showing generation of *bcas2* mutants. Two mutant lines were obtained with mutations that resulted in premature translation termination, resulting in truncated Bcas2 proteins lacking the C-terminal CC domains. (B) Identification of *bcas2* mutations using DNA sequencing. (C) The *bcas2*$^{+/\Delta 7}$ and *bcas2*$^{+/\Delta 14}$ mutants were identified via T7 endonuclease (the upper panel) or FspI restriction enzyme (the lower panel) digestions. (D) Bright field images of embryos derived from crossing indicated females with heterozygous male mutants. Black arrows refer to the embryos that exhibited an abnormal cleavage. The ratio of viable embryos was indicated.

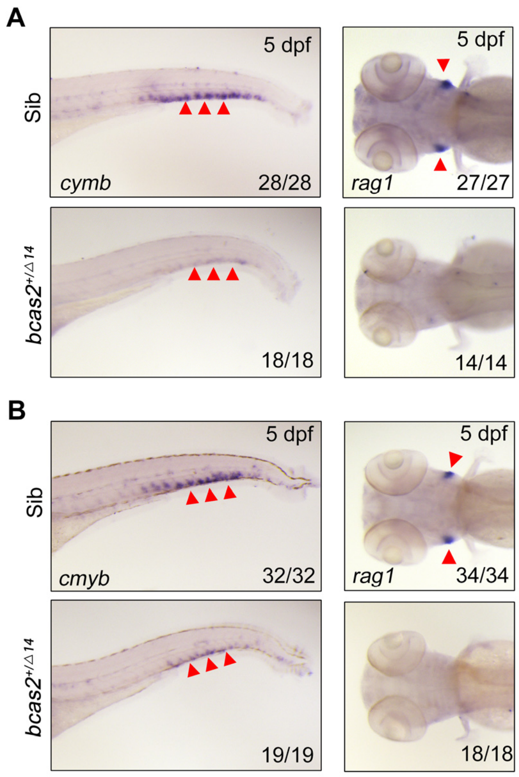


**Figure S3. *bcas2* is essential for definitive hematopoiesis.** (A-B) Expression changes of *cymb* and *rag1* in *bcas2*$^{+/\Delta7}$ (A) and *bcas2*$^{+/\Delta14}$ (B) embryos compared with their siblings. Arrowheads on the left panels indicate the caudal hematopoietic tissue, and those on the right panels indicate the thymus.

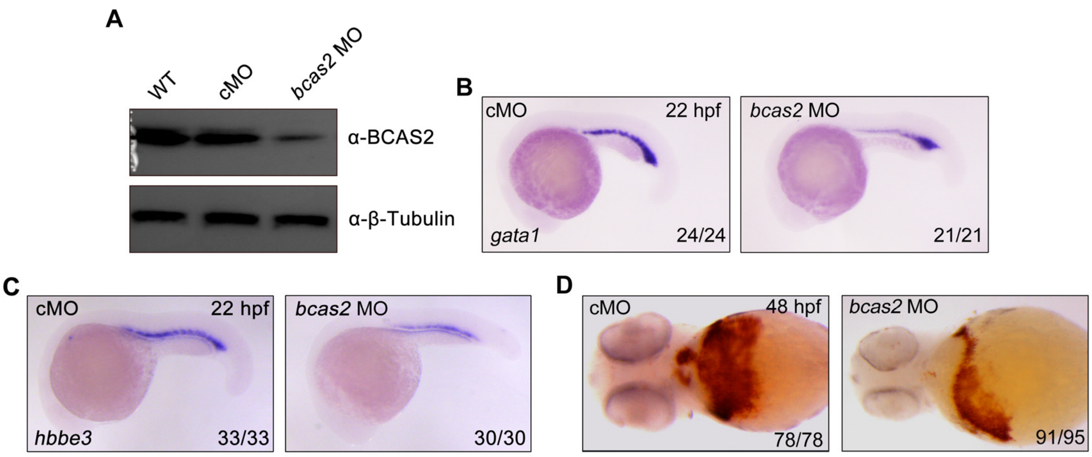


**Figure S4. Knockdown of *bcas2* impairs primitive hematopoiesis.** (A) Western blot analysis showing the expression changes of Bcas2 protein in wild-type embryos and embryos injected with 8 ng control MO (cMO) or *bcas2* MO. (B-C) Expression of *gata1* (B) and *hbbe3* (C) in *bcas2* morphants and control embryos. (D) Detection of hemoglobin levels by *o*-dianisidine staining in *bcas2* morphants and control embryos.

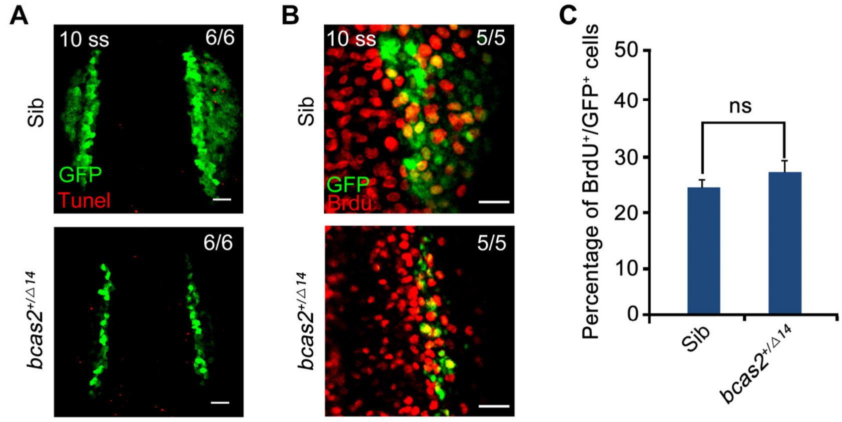


**Figure S5.** ***bcas2*** **is dispensable for the survival and proliferation of hematopoietic cells.** (A) Cell apoptosis was detected by TUNEL assay in *bcas2*$^{+/\Delta14}$ embryos and their siblings with *Tg(gata1:GFP)* background. Scale bar, 50 μm. (B-C) Cell proliferation was detected by BrdU staining in *bcas2*$^{+/\Delta14}$ embryos and their siblings with *Tg(gata1:GFP)* background. Scale bars, 20 μm. The percentage of BrdU$^{+}$/GFP$^{+}$ cells (% over the total number of GFP$^{+}$ cells) was quantified from six embryos in each group (C). Data represent the mean ± SD of three independent experiments. ns, not significant (Student's *t-test*).

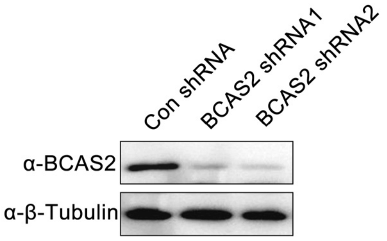


**Figure S6. Western blot analysis of HEK293T Cells transfected with corresponding shRNA constructs.**

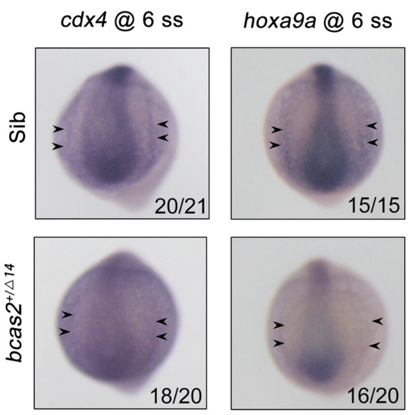


**Figure S7. Expression patterns of *cdx4* and *hoxa9a* in *bcas2*$^{+/\Delta 14}$ embryos and their siblings at the 6-somite stage.** Black arrows indicate the posterior lateral mesoderm.

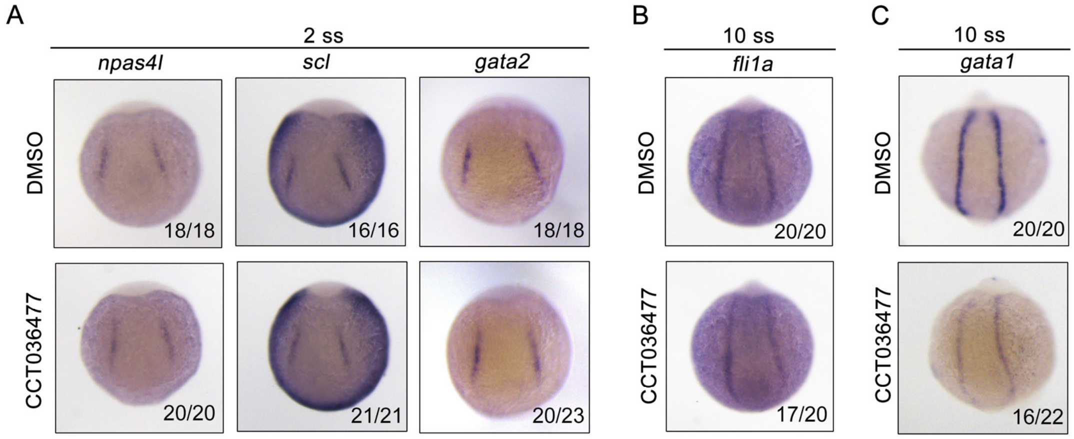


**Figure S8. Inhibition of Wnt signaling does not affect the generation of hemangioblasts or their endothelial differentiation, but impairs their hematopoietic differentiation.** Expression of hemangioblast markers *npas4l*, *scl*, *gata2* (A), endothelial marker *fli1a* (B) and erythroid progenitor marker *gata1* (C) at the indicated stages. Wild-type embryos were treated with 10 μM CCT036477 from 9 hpf and then collected for whole-mount *in situ* hybridizations.

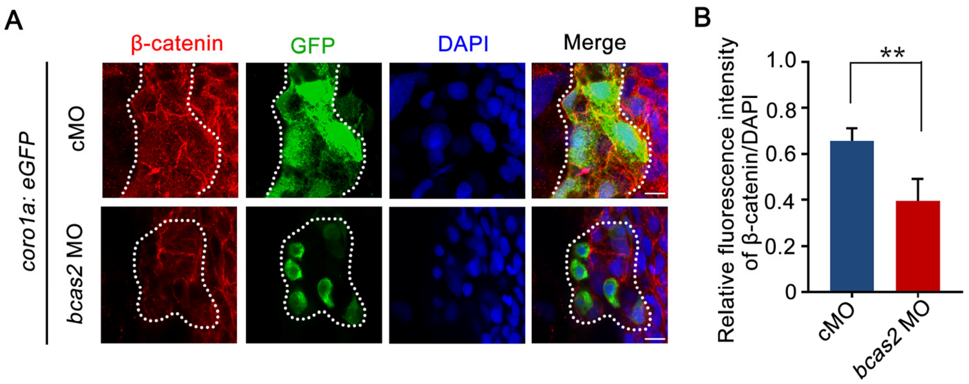


**Figure S9. Knockdown of *bcas2* significantly reduces nuclear β-catenin in the primitive myeloid cells.** Immunofluorescence staining of β-catenin in *Tg(coro1a:GFP)* embryos at 17 hpf. The embryos were injected with 8 ng of the indicated MOs at the one-cell stage and then collected for immunofluorescence staining. The dotted lines refer to the GFP-positive primitive myeloid cells. Scale bars, 10 μm. The relative fluorescence intensity of nuclear β-catenin was quantified in (B). $^{**}P < 0.01$ (Student's *t-test*).

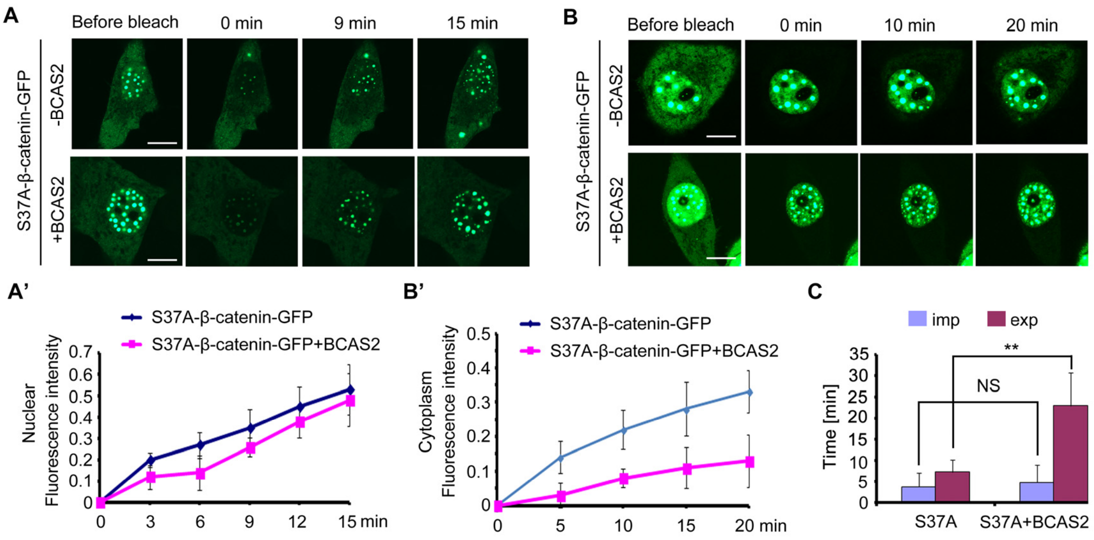


**Figure S10. BCAS2 inhibits the nuclear export of β-catenin.** (A-B') GFP-tagged S37A-β-catenin was co-expressed with or without BCAS2 in HeLa cells. The entire nucleus (A) or entire cytoplasm (B) was bleached. (A-A') Representative images at the indicated timepoints showing the kinetics of nuclear import of S37A-β-catenin are shown in (A) and the related fluorescence recovery curves are shown in (A'). Time-lapse imaging (B) and recovery curves (B') from cytoplasmic photobleaching experiments showed the kinetics of nuclear export of S37A-β-catenin. Scale bars, 10 µm. (C) Quantitative analysis of fluorescence recoveries after photobleaching (n=7). Data represent the mean ± SD of three independent experiments. Student's t-test was used. ns, not significant, $^{**}P < 0.01$ (Student's *t-test*).

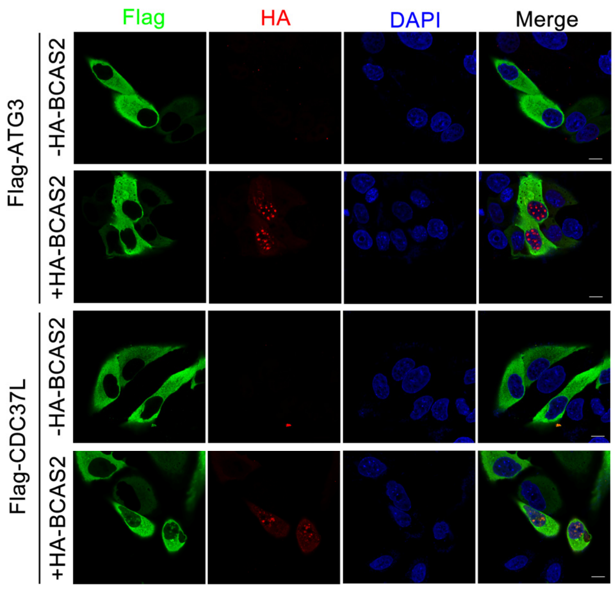


**Figure S11. Overexpression of BCAS2 slightly enhances the nuclear accumulation of CDC37L, and has no influence on the distribution of ATG3.** HeLa cells were transfected with the indicated plasmids and then subjected to immunostaining using anti-Flag and anti-HA antibodies. The nuclei were labeled with DAPI. Scale bar, 10 μm.

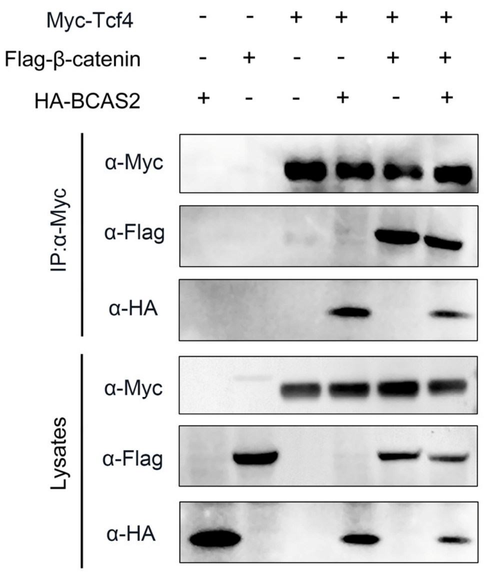


**Figure S12. The interaction between β-catenin and TCF4 remains unaffected in the presence of BCAS2.** HEK293T cells were transfected with the indicated constructs. Cell lysates were immunoprecipitated using anti-c-Myc antibody and the eluted proteins were analyzed by western blotting.

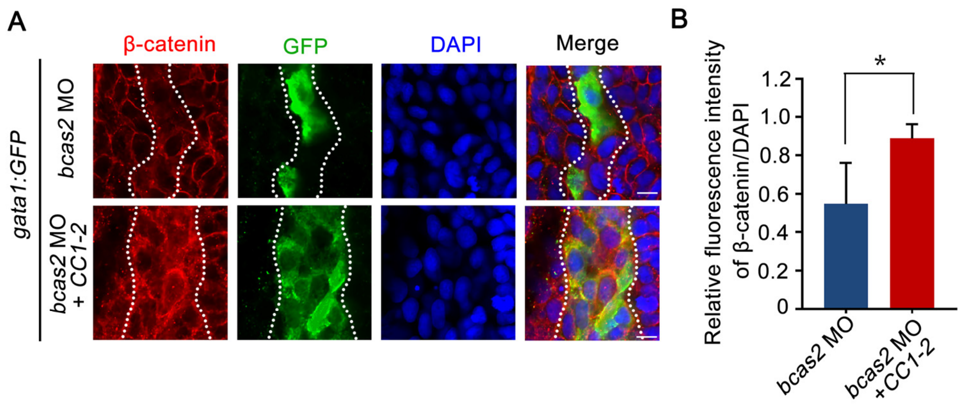


**Figure S13. Overexpression of the CC domains of BCAS2 restores nuclear β-catenin accumulation in *bcas2* morphants.** Immunofluorescence staining of β-catenin in *Tg(gata1:GFP)* embryos at 16 hpf. The embryos were injected with 8 ng *bcas2* MO and 300 pg of *BCAS2 CC1-2* mRNA at the one-cell stage. The dotted lines show the GFP-positive hematopoietic progenitor cells. Scale bars, 10 μm. The relative fluorescence of nuclear β-catenin was quantified in (B). *$P < 0.05$ (Student's *t-test*).

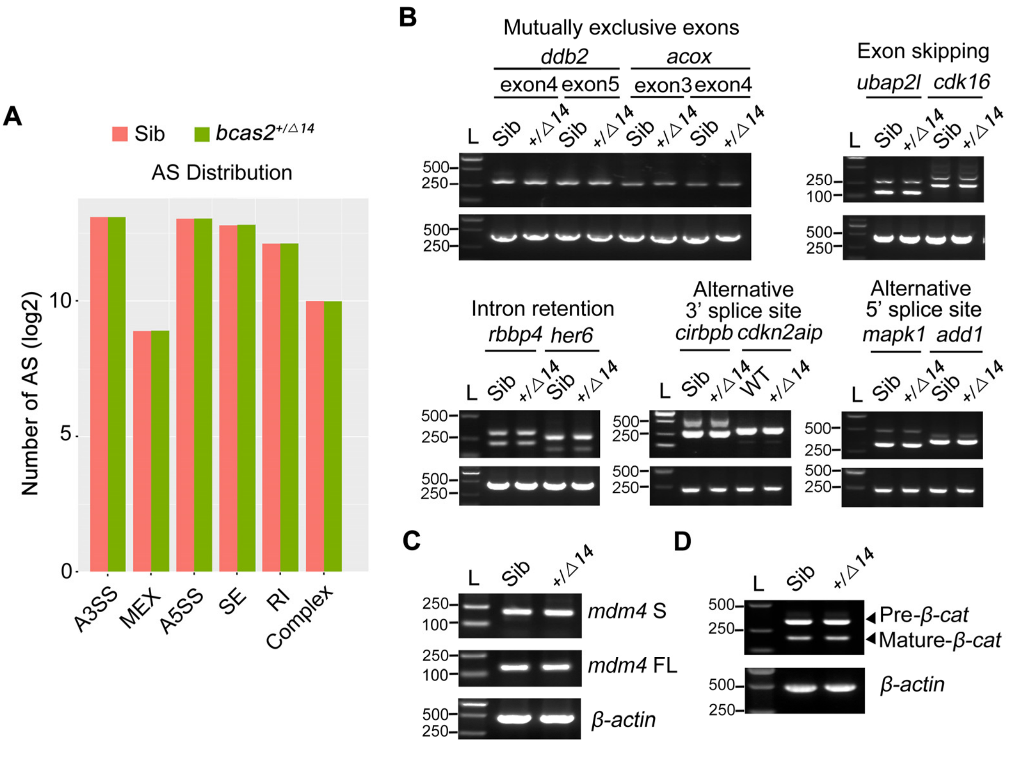


**Figure S14. Haploinsufficiency of *bcas2* does not affect pre-mRNA splicing during primitive hematopoiesis.** (A) The number of five major types of alternative splicing events was analyzed from RNA sequencing data. Embryos were lysed at the 10-somite stage and subjected to reverse transcription. The cDNA library was prepared, sequenced and then analyzed using rMATS. The five main alternative splicing types refer to exon skipping (SE), alternative 3' splicing site (A3SS), alternative 5' splicing site (A5SS), intron retention (RI), and mutually exclusive exons (MEX). (B) Examples of different types of alternative splicing were analyzed by reverse transcription PCR using total RNA from sibling and *bcas2*$^{+/Δ14}$ embryos at the 10-somite stage. (C) Reverse transcription PCR analysis of total *mdm4*-FL and *mdm4*-S isoforms in *bcas2*$^{+/Δ14}$ and sibling embryos at the 10-somite stage. (D) Analysis the pre-mRNA and mature mRNA of *β-catenin* in $bcas2^{+/\Delta 14}$ and sibling embryos at the 10-somite stage.
